# Neural differential equations enable early-stage prediction of preterm birth using vaginal microbiota

**DOI:** 10.1101/2023.09.22.558954

**Authors:** Kaushik Karambelkar, Mayank Baranwal

## Abstract

Preterm births (PTBs), i.e., births before 37 weeks of gestation are completed, are one of the leading issues concerning infant health, and is a problem that plagues all parts of the world. Millions of infants are born preterm globally each year, resulting in developmental disorders in infants and increase in neonatal mortality. Although there are known risk factors for PTB, the current procedures used to assess PTB risk are effective only at the later stages of pregnancy, which reduces the impact of currently possible interventions administered to prevent PTB or mitigate its ill-effects. Vaginal microbial communities have recently garnered attention in the context of PTB, with the notion that a highly diverse microbiome is detrimental as far as PTB is concerned. Increased abundance or scarcity of certain microbial species belonging to specific genera has also been linked to PTB risk. Consequently, attempts have been made towards establishing a correlation between alpha-diversity indices associated with vaginal microbial communities, and PTB. However, the vaginal microbiome varies greatly from individual to individual, and this variation is more pronounced in racially, ethnically and geographically diverse populations, which diversity indices may not be able to overcome. Machine learning (ML)-based approaches have also previously been explored, however, the success of these approaches reported thus far has been limited. Additionally, microbial communities have been reported to evolve during the duration of the pregnancy, and capturing such a signature may require higher, more complex modeling paradigms. Thus, alternative approaches are necessary to identify signatures in these microbial communities that are capable of distinguishing PTB from a full-term pregnancy. In this study, we have highlighted the limitations of diversity indices for prediction of PTB in racially diverse cohorts. We applied Deep Learning (DL)-based methods to vaginal microbial abundance profiles obtained at various stages of pregnancy, and Neural Controlled Differential Equations (CDEs) are able to identify a signature in the temporally-evolving vaginal microbiome during trimester 2 and can predict incidences of PTB (mean test set ROC-AUC = 0.81, accuracy = 75%, F1-score = 0.71) significantly better than traditional ML classifiers such as Random Forests (mean test set ROC-AUC = 0.65, accuracy = 66%, F1-score = 0.42) and Decision Trees (mean test set ROC-AUC = 0.48, accuracy = 46%, F1-score = 0.40), thus enabling effective early-stage PTB risk assessment.

**Graphical Abstract:** 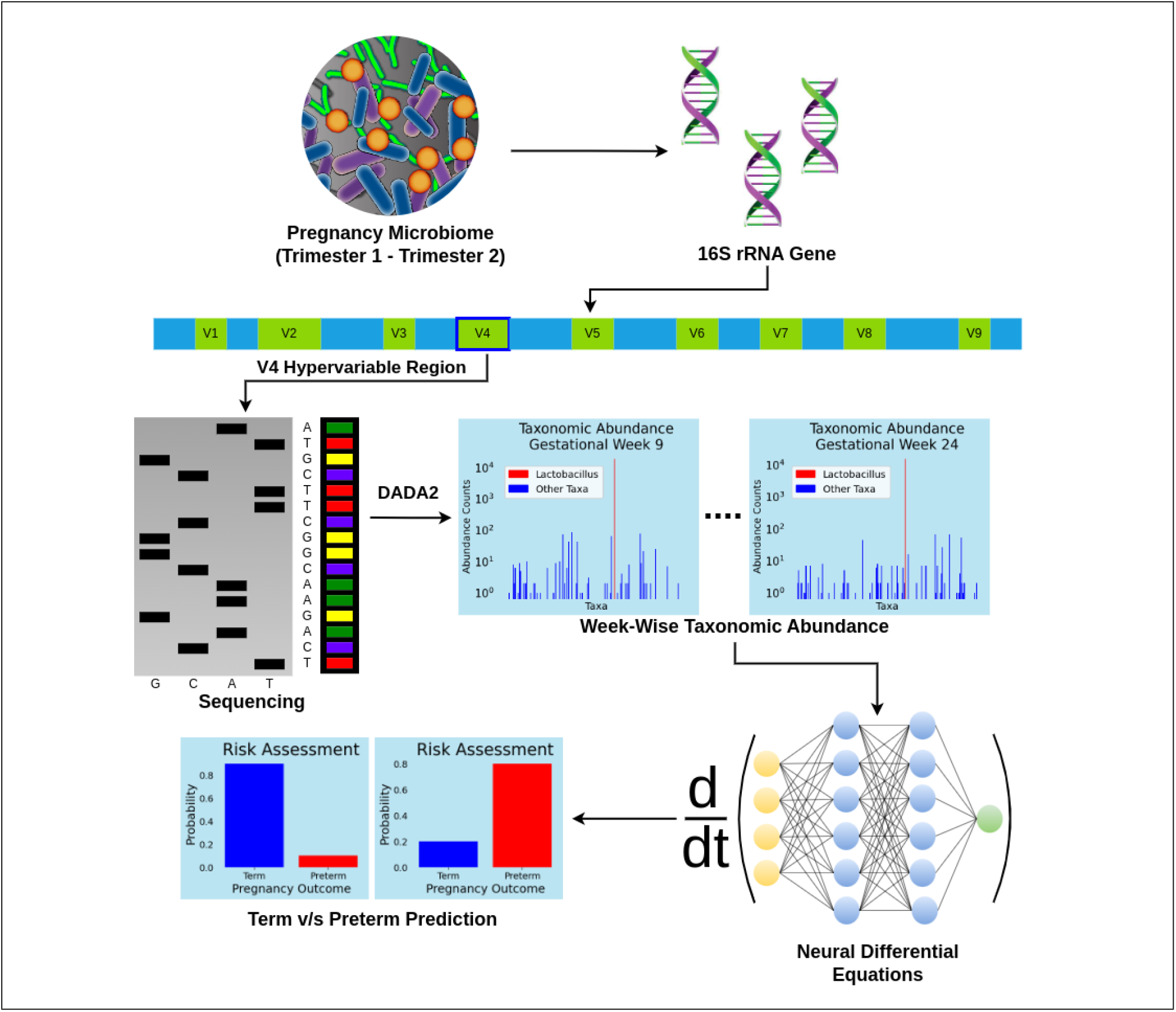

## 1 Introduction

Preterm births (PTBs) are live births that occur before 37 weeks of pregnancy and are a major public health concern worldwide. It is estimated that about 15 million babies are born pre-term globally each year, putting the global PTB rate at about 11% [8]. PTB is among the leading causes of neonatal mortality and morbidity, especially in low- and middle-income economies [52]. The global PTB rate is also on a steady rise, thus making it a significant global burden [14]. Approximately 18% of the deaths among children under the age of 5 years happen within the first 28 days of life and can be attributed to complications arising from PTB [63]. Additionally, it can lead to long-term health complications such as respiratory illnesses, neurodevelopmental disorders and learning disabilities, arising from the developmental issues associated with PTB [62, 16]. PTB is also not a problem that is specific to underdeveloped or developing countries, although the ill-effects of it may be more pronounced in low-income countries [6, 12]. Incidences of PTB are also found in high-income parts of the world, albeit with a lower frequency, and the rates of PTB have not been on the decline either in most parts of the world [14].

The pathophysiology behind PTB is not completely understood yet, although certain risk factors, including but not limited to smoking habits, alcohol intake and reproductive history have been identified to be associated with increased pre-term delivery risk [8, 53, 59]. The ill-effects of PTB can be mitigated, and a healthy, full-term gestation outcome may also be achieved if appropriate interventions are administered [32, 45]. Their success, however, depends on identifying at-risk subjects at earlier stages of their pregnancy, as these approaches are effective when administered at earlier stages of pregnancy [8, 45]. Current methods for assessing pre-term pregnancy outcomes involve the use of physical and biochemical markers, which are not accurately determinative of a potential incidence of premature culmination of pregnancy in the future [13, 25].

Many non-pathogenic bacteria, viruses and fungi inhabit various areas of the human body, such as the gut, mouth, and reproductive tracts [56], and are collectively referred to as the “microbiome”. Microbiomes are essential for normal functioning of the respective organs, and maintain a symbiotic relationship with the human body and drive key biochemical reactions, [27, 33] and dysregulated microbiomes are often implicated in various diseases. These microbial communities are also present in the reproductive tracts, and have been reported to influence the pregnancy outcome [41]. There is evidence linking the composition of vaginal microbiomes to risk of PTB, and the abundance levels of specific microbiota, such as various species of the *Lactobacillus* genus, have the potential to be indicative of PTB even at earlier stages of pregnancy [37, 17, 54, 9]. Vaginal microbial communities can be categorized into specific Community State Types (CSTs), which are typically characterized by abundances of various *Lactobacillus* species [54]. CSTs are associated with increased or decreased risk of abnormalities such as Bacterial Vaginosis (BV), Urinary Tract Infections (UTIs) and even PTB [28]. Moreover, alpha-diversity indices, such as Shannon and Simpson diversity, which can quantify the diversity of vaginal microbiota, have been harnessed for predicting PTB [18, 34, 31]. However, the vaginal microbiome differs considerably from individual to individual, especially across races [60, 30]. Additionally, the microbial abundance may further vary depending on the sequence processing methods used on 16S ribosomal RNA (rRNA) data, which is typically used to estimate taxonomic abundance at various levels of classification [7]. Consequently, the success of diversity indices for estimating PTB risk may be specific to certain cohorts, or be influenced by the sequence processing pipeline and consequently, may not translate across cohorts, as is our observation in this study. Machine Learning (ML)-based approaches have also been explored in this context, which leverage features such as abundance of various taxa, phylotype counts, CST of the vaginal microbiome, age, race and more for PTB risk assessment. The Dialogue for Reverse Engineering Assessments and Methods (DREAM) Preterm Birth Microbiome Prediction Challenge was launched recently, incentivizing ML approaches for PTB risk assessment using the vaginal microbiome. The challenge consisted of two sub-challenges: one for term vs preterm prediction (birth after vs before 37 weeks of gestation) and another for early preterm prediction (birth after vs before 32 weeks of gestation). The best submissions in the first sub-challenge involved the use of tree-based ML methods, but the quality of the predictions, even for the top-performing models, was not sufficient for clinical translation.

The vaginal microbiome evolves as the pregnancy progresses [41, 54, 64], and the numerous changes that it undergoes may contain a signature for identifying PTB risk. Currently, there is a severe lack of approaches that exploit the temporal dynamics of vaginal microbiomes for PTB risk assessment. PTB is also categorized according to the extent of prematurity, namely extremely preterm (< 28 weeks), very preterm (28-32 weeks) and moderate to late preterm (32-37 weeks) [4], each having varying impact on infant health [4, 58]. Current approaches exploiting microbiome data also lack the capability to estimate the period of delivery in cases of PTB, which is very important when administering interventions to attempt to achieve a full-term pregnancy or mitigate the adverse outcomes of PTB.

Deep learning approaches such as Recurrent Neural Networks (RNNs), have previously been used for modelling the dynamics of gut and other microbiota in various contexts [3, 24, 43] and have found success. To the best of our knowledge, such methods have not been applied to vaginal microbiota, especially in the context of PTB risk assessment, so far. This may be partly attributed to the fact that RNN-based approaches demand data sampled at regular intervals, which is challenging to collect as study subjects are often irregular in clinical visits. With this study, we present a deep learning-based approach that is capable of differentiating between term and preterm births using temporal vaginal microbiome data, which overcomes the dependence on regularly sampled microbial data. We also highlight the limitations of alpha diversity indices and traditional ML methods for PTB prediction in racially and ethnically diverse patient cohorts. We show that modeling the temporal dynamics of microbiota using deep learning methods results in more reliable PTB risk scoring than simple ML-based methods. Our best model, utilizing Neural Controlled Differential Equations (CDEs), predicts PTB with substantial accuracy (mean test set ROC-AUC = 0.82, accuracy = 75%, sensitivity = 0.71, specificity = 0.85, F1-score = 0.71), thus outperforming any ML-based PTB prediction approaches so far. On the basis of this study, we show the potential of vaginal microbiota for PTB prediction, and that such approaches can be pushed towards complete clinical viability with further efforts.

## 2 Methods

### 2.1 Dataset

We obtained 16S rRNA sequences collected from human patient samples. The data was sourced from a previously published study by Callahan et al. on refinement of a vaginal microbiome signature of preterm birth [10]. The dataset is publicly available under the open access category in the Sequence Read Archive (SRA), BioProject ID PRJNA393472. It consists of 16S rRNA sequence samples, spread across 133 racially and ethnically diverse subjects, and sampled at different points of time during the pregnancy.

### 2.2 16S rRNA Sequence Processing

It has been widely established that the hypervariable regions (V1-V9) within 16S rRNA gene can be used for phylogenetic studies and genus or species-level classification in diverse microbial populations [65]. Furthermore, certain hypervariable regions (such as the V4) are semi-conserved and can reliably predict specific taxonomic levels [66]. The procedure to convert the 16S rRNA sequence data to microbial abundance involves various stages of processing. In the first step, quality control checks are performed and sequencing artifacts, low quality reads, etc. are removed from the read sequences. Secondly, the preprocessed sequences are aligned against a chosen reference database, and a taxonomic class is assigned to each sequence. The sequences are then grouped into clusters, which represent Operational Taxonomic Units (OTUs), based on sequence similarities. Lastly, the abundances of each OTU in samples are estimated [55, 20, 21, 11]. The microbial abundance obtained depends on the specific processing steps, and variations in processing steps can result in different abundance values [55, 20, 21, 11]. We used the DADA2 processing pipeline [11] to derive microbial abundance data from the sequence reads. The metadata and taxonomic abundance tables were generated using the SRA cloud [35] and abundances were obtained at various levels of taxonomic classification. We retained genus-level abundances for all our analyses since the resolution at which lower-level abundances were captured was much lower.

### 2.3 Processing Taxonomic Abundance Data

We eliminated the samples for which metadata information for certain key fields, viz., gestational age at the time of sample collection, gestational age at delivery, etc., was missing. We eliminated the genera abundance for samples collected during trimester 3 (gestational age > 24 weeks) for our analyses, with the intention of being able to predict instances of preterm delivery sufficiently early. Furthermore, we removed samples collected during or before the 8^th^ week of gestation, as they were present for very few (6 out of 133) subjects. Furthermore, for some of the analyses, we transformed the genera abundance data to sample-wise relative abundance, and filtered out genera with high skewness (> 10 and < −10) and high kurtosis (> 10 and < −10). We retained 70% of the subjects (90 out of 133) as the training dataset and the rest were used for validating the approaches. The training and test datasets were kept consistent across all the analyses.

### 2.4 Diversity Metrics

Alpha diversity metrics have been reported to be potentially indicative of preterm birth, and a highly-diverse vaginal microbiome is correlated with increased risk of preterm delivery [18, 34, 31]. We computed Shannon, Simpson, Chao1 and Gini alpha diversity indices, as well as Taxonomic Composition Skew (TCS) [31], a diversity index specifically tailored for the vaginal microbiome. Unlike other diversity indices, TCS takes into account that vaginal microbiomes are usually dominated by the *Lactobacillus* species and other genera are in the minority. TCS responds in a different manner, to changes in abundances of sparse and dominant taxa, and thus is possibly more suitable for quantifying the diversity in vaginal microbiomes. Based on the computed values of these metrics, we searched for an optimal threshold to differentiate between term/preterm delivery using the training set, and validated it on the test set. The standard diversity metrics were computed using the `scikit-bio` python library (version 0.5.8).

### 2.5 Traditional ML Approaches

We used two ML classifiers: Decision Tree (DT) and Random Forest (RF), to predict term/preterm outcomes. This constituted a secondary baseline for benchmarking the performance of higher, more complex deep learning-based prediction approaches. For each patient subject, the microbial abundance profile closest to the week of delivery and obtained during the period between the 9^th^ and the 24^th^ week of gestation, following the hypothesis that composition of vaginal microbial communities closer to the period of delivery are better indicative of preterm delivery risk. The resultant training and test sets contained 93 and 40 samples respectively.

We carried out 3-fold cross-validation on the training set to optimize the hyperparameters of each classifier by performing a grid search on a pre-defined search space for each parameter. For decision tree, we optimized the maximum depth of the tree (`max_depth`), number of leaf nodes to grow the tree (`max_leaf_nodes`), minimum number of samples required to be at a leaf node (`min_samples_leaf`), minimum number of samples required to split an internal node (`min_samples_split`), number of variables to consider while selecting best split (`max_features`), as well as the splitting strategy, loss function and the weight ratio to account for class imbalance. For random forests, we optimized the number of trees (`n_estimators`), `max_depth`, `max_leaf_nodes`, `min_samples_leaf`, `min_samples_split`, loss function and the class weight ratio. The search spaces for the respective parameters are listed in supplementary table 1. We assessed the predictive performance of these classifiers on the test set using validation metrics such as area under the receiver operating characteristic curve (ROC-AUC), accuracy, precision, recall and F1-score. Implementations for both classifiers were derived from the scikit-learn python library (version 1.2.2).

### 2.6 Deep Learning Approaches

Machine learning classifiers, such as Support Vector Classifiers (SVCs), as well as tree-based classifiers such as DT and RF which we explored as well, have been explored extensively for preterm birth prediction using vaginal microbiota, most often in tandem with other features such as physical markers and patient history. However, these classifiers have largely lacked the capability of making reliable predictions. Surprisingly, deep learning models have hardly been explored for this particular problem. Given the time-series nature of the data, we focused on deep learning algorithms for sequential data in this study.

#### 2.6.1 Recurrent Neural Networks

Recurrent Neural Networks (RNNs) are a type of neural networks designed to handle sequential or time-series data. The issue with standard RNNs however, is that they have difficulty in learning long-term dependencies in long sequences, due to the issue of vanishing/exploding gradients [50]. Long Short-Term Memory (LSTM) is a type of RNN that is capable of learning long sequences, and are possibly more appropriate for the week-wise taxonomic abundance dataset. LSTM maintains a hidden state, which stores short-term information, and a cell state, which stores long-term information. The initial hidden and cell states are generally set to zero vectors. A LSTM cell at each time step updates the hidden and cell states based on the states at the previous time step and the input data at the current time step.

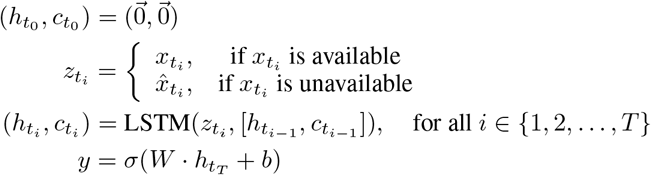

However, the conventional LSTM system demands a continuous and uniform time-series dataset, i.e., the time-steps must represent uniform intervals and the input data should be available for each step. While the taxonomic abundance data is uniformly sampled (week-wise), data for some subjects is not present for some weeks. Additionally, the first week for which data is available is different for each subject, and thus, a zero vector initialization will not be appropriate for the initial hidden state. To address these issues we modified the LSTM network accordingly. Firstly, we initialized the hidden state 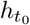 with a trainable embedding layer, and the cell state 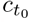 was initialized as a zero vector. Secondly, at each time step, we had the LSTM cell generate the taxonomic abundance forecast 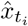 at each time step, and used the forecast whenever the input data 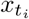 was not available. The final hidden state 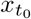 was fed to a linear layer with parameters *W* and *b* followed by a sigmoid activation *σ*(·) to predict the term/pre-term outcome (*y*). The entire network was trained end-to-end.

Figure 1 outlines the described LSTM network. The LSTM implementation assumes that the input data is continuously and regularly sampled. However, in our case, data for some intervals may be missing. To overcome this, we masked the data for missing time steps by using zero vectors to substitute the missing data points. In parallel, we also used a vector indicating the coordinates of the masked time intervals for each sample, for which the model used the forecast, i.e., the model-predicted microbial abundance instead of the ground truth values. Additional method details are provided in the supplementary material (section 2.1) and the hyperparameter values are listed in supplementary table 3. The LSTM model was implmented using the pytorch library [51] (version 2.0.1, cuda version 11.8).

**Figure 1:**
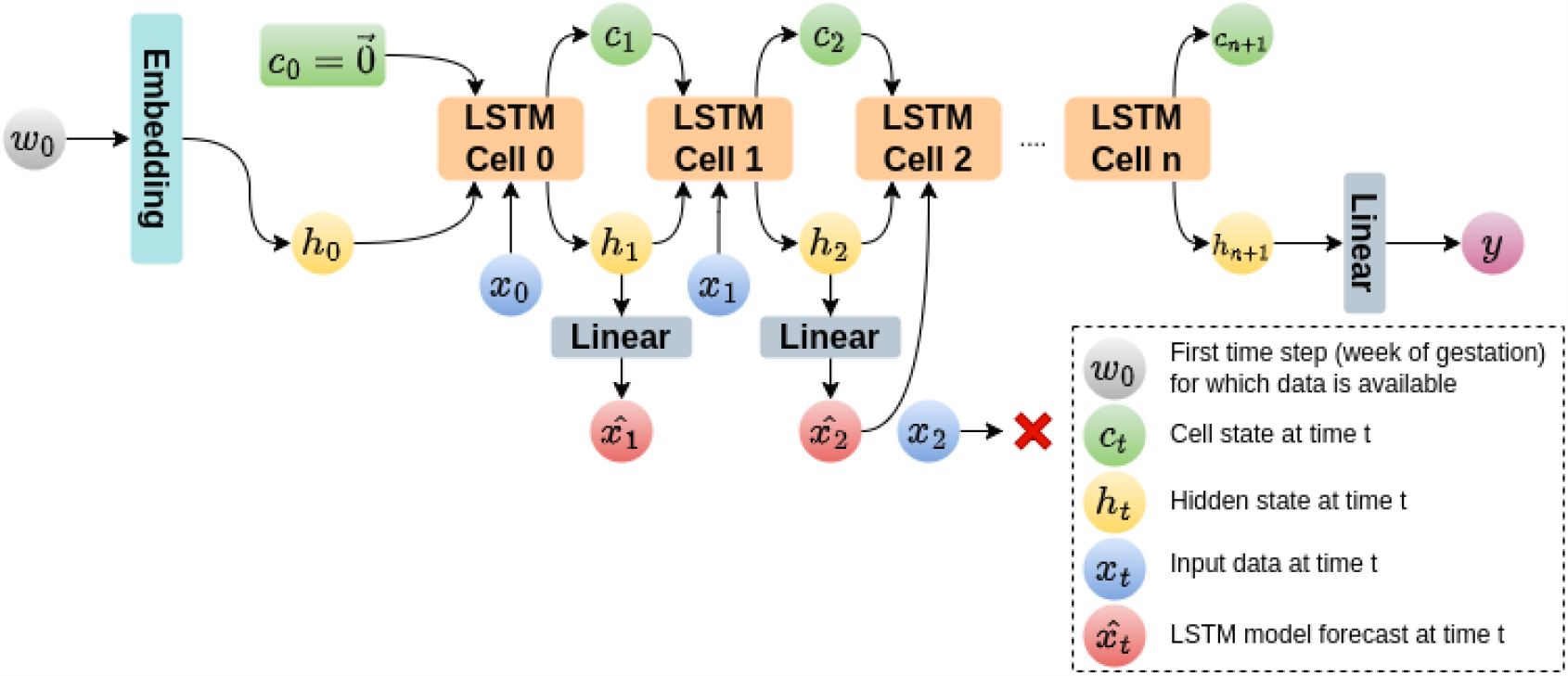
Visualization of the LSTM Network. Week 0 represents the earliest week of gestation for which microbial data is available. This is passed through an embedding layer, which generates the initial hidden state. Subsequent hidden states are determined by the previous hidden and cell states, and the microbial data at the respective time step. If the microbial data is not available, the output of the previous hidden state is used instead. The final hidden state is passed through a linear layer with a single output neuron to predict the term/preterm outcome

#### 2.6.2 Neural Differential Equations

A significant limitation associated with clinical data pertains to its irregular sampling of data points, which presents challenges in constructing effective machine learning models that can effectively harness the inherent time-series information. The irregularity in data sampling introduces two notable drawbacks: firstly, the size of input data, contingent upon the number of sampling instances, differs among various subjects; secondly, the timing of sampling instances is not strictly discrete, thereby restricting the applicability of commonly employed RNN models that assume uniform intervals between sampled data points. To overcome this, we leverage a recently introduced class of deep learning models - Neural Ordinary Differential Equations (ODEs) that combine a neural network with ODEs and allow for continuous interpolation between two randomly spaced sampling instants.

Notably, Neural ODEs exclusively consider the evolution of time-series data commencing at a fixed time point, denoted as *t*_0_, which is accompanied by an initial condition represented as 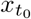. In the context of our research, 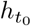signifies the initial abundance of genera at this specific time *t*_0_, which might correspond to the onset of gestation week 9. Regrettably, the trajectory of taxa abundance varies from person to person, and the initial abundance data for all subjects may not be accessible for subsequent analysis, as the microbial profiles of each subject were not uniformly sampled at the same time point, namely *t*_0_. Consequently, while Neural ODEs excel in interpolation tasks, they cannot be seamlessly integrated into our framework due to the potential absence of consistent initial abundance data. Nevertheless, it transpires that addressing this problem, specifically how to integrate incoming information, has already been thoroughly explored within the realm of mathematics, particularly in the field of rough analysis, which is dedicated to the examination of *controlled differential equations* (CDEs) [39, 40]. Kidger et al. [36] have introduced a novel framework known as Neural CDEs, which extends CDEs to Neural ODE models. To put it simply, Neural CDEs can be seen as continuous-time counterparts of Recurrent Neural Network (RNN) models. These models can be trained efficiently using a method called “adjoint backpropagation”, which is elaborated on briefly in the supplementary materials (section 3.1), and detailed mathematical representation of it can be found in [15]. In brief, the Neural CDE model can be summarized through the following sequence of operations:

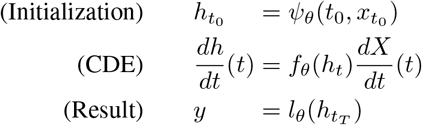

Here *ψ*_*θ*_ and *l*_*θ*_ correspond to linear models responsible for transforming the initial taxa abundance (along with the time-stamp *t*_0_) into the initial hidden state 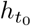 and the final hidden state 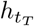 into the output label, respectively. The map *ψ*_*θ*_ is used to avoid translational invariance to first sampled time instant. *X* is the natural cubic spline with knots at *t*_0_, …, *t*_*T*_ such that 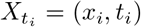. Natural cubic splines allow for smooth interpolation and minimum regularity for handling certain edge cases. *f*_*θ*_ is neural network model depending on parameters *θ*. Due to its dependence on cubic splines, Neural CDEs [36] can be applied to irregularly sampled time series, even with temporally-scattered initial conditions. Thus, we chose to apply Neural CDEs to predicting preterm birth using the irregularly-sampled microbial abundance dataset. A comprehensive mathematical introduction to Neural CDEs is outside the scope of this paper. For those interested in delving deeper into the mathematical details, we recommend consulting [36] for a more thorough explanation. The torchcde library (version 0.2.5) was used to implement the Neural CDE model in python. Hyperparameter values for the model are listed in supplementary table 4.

## 3 Results

### 3.1 Microbial abundance dataset contains racially and ethnically diverse subjects

The 16S rRNA sequence data was converted to taxonomic abundance (Methods, section 2.2), which led to approximately 290,000 abundance counts, spanning taxa counts at various levels of classification, out of which approximately 65,000 corresponded to genus-level, belonging to 2,326 unique samples which collected at various weeks of gestation throughout the pregnancy, spread across 133 subjects of diverse race and ethnicities. We aligned the abundance counts subject- and gestational week-wise for ease of interpretation. Abundance counts for multiple samples derived from the same subject collected during the same week of gestation, if any, were replaced by the mean of those counts to ensure consistency, as future analyses were carried out on week-wise data. Figure 2 describes various trends in the microbial abundance data. Microbial abundance profiles corresponding to 43 out of the 133 subjects were reserved as the test set. (refer methods section 2.3).

**Figure 2:**
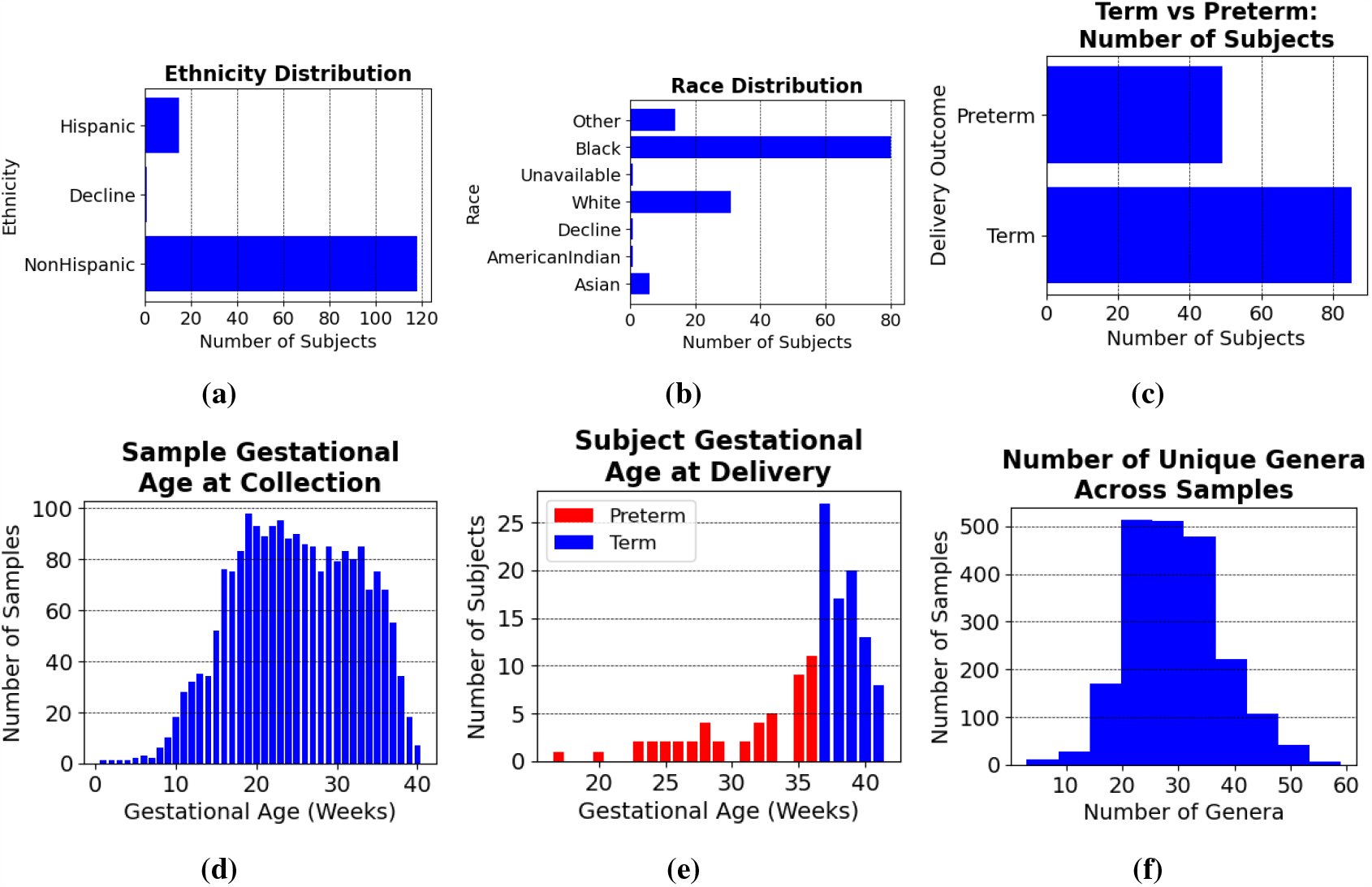
Distribution of a Ethnicity, b Race, c Number of subjects who delivered at term/pre-term, d Sample gestational age at the time of collection, e Gestational age of subject at the time of delivery f Number of unique genera detected in each sample, in the microbial abundance dataset.

### 3.2 Diversity metrics do not reliably identify at-risk PTB subjects

Alpha-diversity indices were computed on samples collected during trimester 1 and trimester 2 (gestational weeks 9 to 24, see methods section 2.3). The visualization of alpha-diversity metrics computed at various weeks of gestation, along with the delivery outcome (term/pre-term) is presented in figure 2.4. No signature that can distinguish term/preterm birth is observable from these metrics. As expected from the plot, the alpha-diversity indices are not predictive of a preterm delivery outcome, and perform worse than random classifiers (prediction accuracy on test set < 50%, refer methods section 2.4).

**Figure 3:**
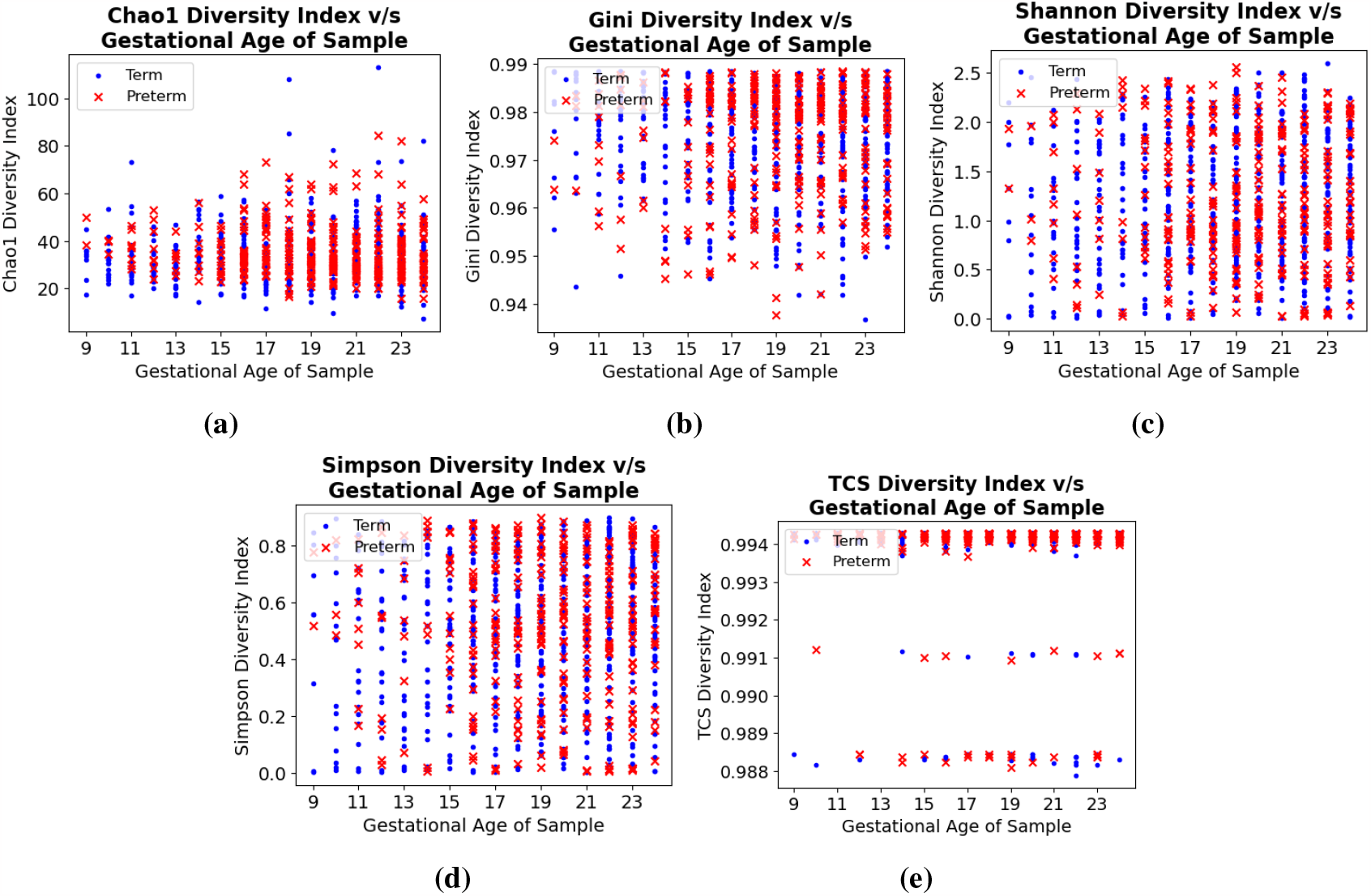
a Chao1, b Gini, c Shannon, d Simpson and e TCS diversity metrics computed on microbial abundance data collected during trimester 1 and trimester 2 of pregnancy. Blue and red points represent samples derived from subjects who delivered at term and pre-term, respectively.

**Figure 4:**
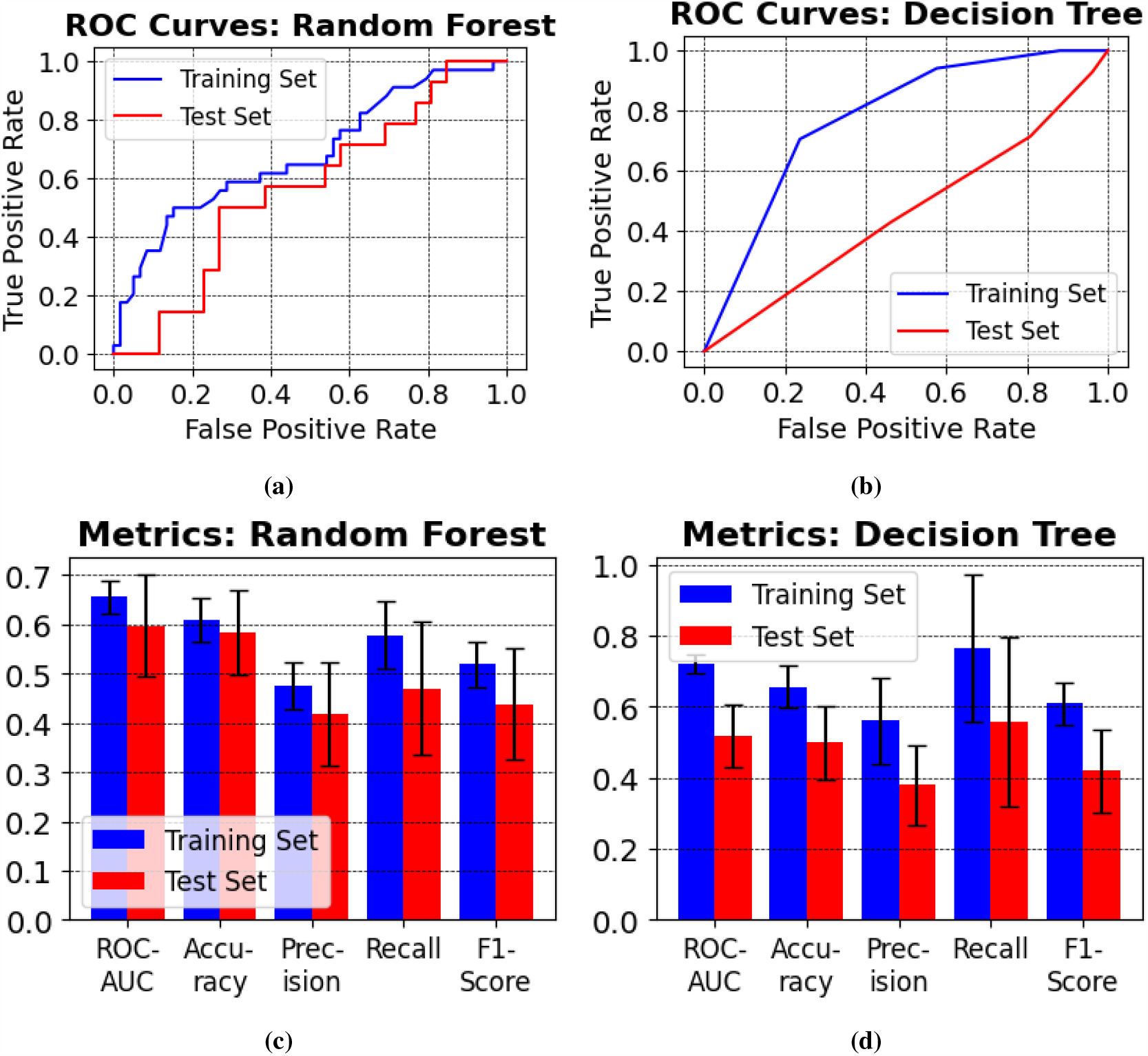
Receiver operating characteristic (ROC) curve for the a random forest and b decision tree classifiers on the training and test datasets, and validation metrics (AUC, accuracy, precision, recall and f1-score) computed for the c RF and d DT classifiers. The error bars in figures c and d account for the stochasticity of the classifiers.

### 3.3 ML classifiers do not make adequate PTB risk assessment

We tested the performance of two ML classifiers, viz., RF and DT towards prediction of preterm birth. For this, we isolated the microbial abundance profiles of training set subjects as well as the test set subjects, collected during the period between the 9^th^ and the 24^th^ week of gestation, thus formulating the training and the testing datasets, respectively. Optimal hyperparameters for both RF and DT were identified by performing a grid search on pre-defined parameter search spaces using 3-fold cross-validation on the training set, and are listed in supplementary table 2. The resultant models were validated on the test set by computing ROC-AUC, accuracy, precision-recall and F1-score. The RF model performed significantly better (mean test set ROC-AUC = 0.60; accuracy = 58.7%, precision = 0.42, recall = 0.48, F1-score = 0.44) compared to DT (mean test set ROC-AUC = 0.46; accuracy = 45.5%, precision = 0.39, recall = 0.58, F1-score = 0.41) which performs worse than a random predictors. Neither of the models, however, make adequately reliable predictions on the test set. The full results for both models are presented in figure 4.

### 3.4 Statistical analyses indicate presence of signatures for PTB prediction in evolving microbiomes

Skewness and kurtosis were computed on relative abundance of microbial genera during the period of trimester 1 - trimester 2 (gestational weeks 9 to 24). The results are presented in figure 5 a, b. We excluded the genera which had highly positively and negatively skewed relative abundances (i.e., skewness > 10 and skewness < −10) as well as genera with high positive and negative kurtosis (kurtosis > 10 and kurtosis < −10). As a result, only 6 genera were retained, namely, *Lactobacillus, Anaerococcus, Gardnerella, Peptoniphilus, Finegoldia* and *Prevotella*. Except for *Finegoldia*, these genera have been identified to be linked to PTB risk previously. *Lactobacillus* is the most dominant genus within the vaginal microbiome, and low counts of *Lactobacillus* have previously been stated to be indicative of increased PTB risk [5, 28]. Certain species of *Anaerococcus* are found to be associated with increased PTB risk [1], however, there are also reports that *Anaerococcus* may be a protective taxon against PTB [19]. There is strong evidence linking high *Gardnerella vaginalis* presence with PTB and bacterial vaginosis, which also increases PTB risk [44, 46]. In some populations, increased counts of certain *Peptoniphilus* species were also found to be associated with high PTB risk [47]. Similar evidence exists associating high *Prevotella* abundances with increased PTB risk [23, 22, 48]. However, there is insufficient evidence to establish a link between these observations and the changes that vaginal microbiota undergo throughout the duration of the pregnancy.

**Figure 5:**
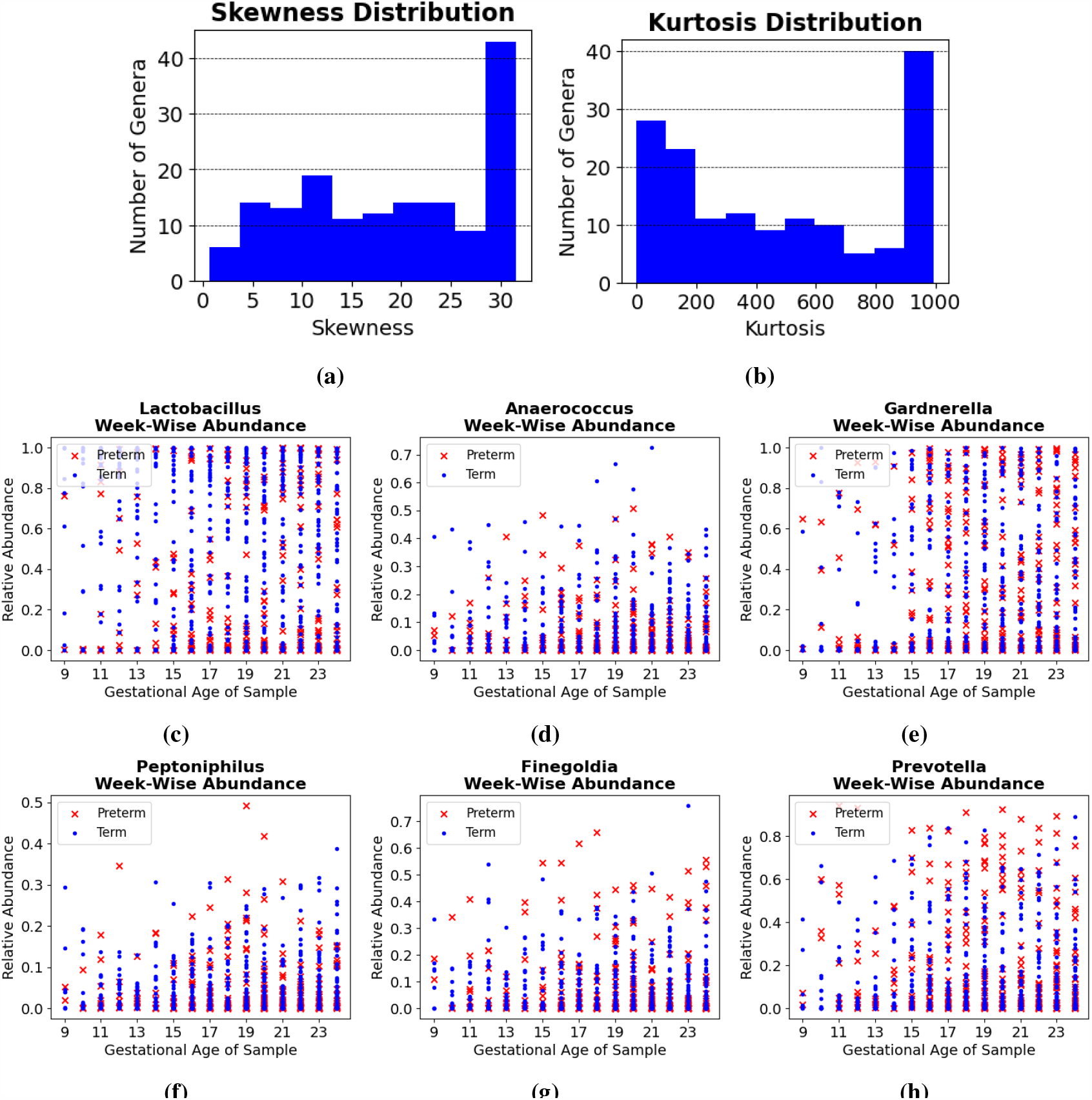
a Skewness and b Kurtosis distribution computed on the relative abundance of each microbial genus during the period from gestational weeks 9 to 24. c -h show changing relative abundances for the Lactobacillus, Anaerococcus, Gardnerella, Peptoniphilus, Finegoldia and Prevotella genera respectively, during the same period.

We further re-computed the relative abundance based on the abundance counts of these 6 genera only, and visualized the gestational week-wise abundances and corresponding term/preterm delivery outcomes. The results are presented in figure 5 c -h. This analysis highlights that the composition of the 6 genera mentioned above, changes significantly during the period from end of trimester 1 to trimester 2 of pregnancy. The trends indicate that low abundance of the *Lactobacillus* genus during trimester 2 (gestational week 16 onwards) may be correlated with increased PTB risk (see figure 5 c), and is consistent with current knowledge. Additionally, we also observed the association of increased *Gardnerella* counts with PTB risk (see figure 5 e), and the effect is more pronounced during gestational weeks 16-18. Most of the samples with increased *Prevotella* counts during weeks 19-21 belonged to subjects who went on to deliver preterm (see figure 5 h).

### 3.5 LSTMs do not decode the temporal dynamics of vaginal microbiomes

LSTM was trained on the week-wise genera abundance data sampled during gestational weeks 9 to 24, after making appropriate adjustments to account for the irregularity in sampling (see Methods section 2.6.1). When trained on the entire set of genera with non-zero variance in the training set, the model overfits and does not generalize well to making predictions on the test set (training set accuracy = 100%, test set accuracy < 60%). Even when trained on the set of genera with low skewness and kurtosis, the model fails to make sufficiently accurate predictions on the test set (accuracy = 63%), and is outperformed by the RF model described above. We attribute this lack of predictivity to the increased estimations that the model makes to fill the temporal gaps in the data, and it may necessitate availability of additional data samples to be able to make these estimations more accurately, either in terms of more patient subject or increased density of samples per subject.

### 3.6 Neural CDEs are capable of achieving PTB prediction with a substantial accuracy

The neural CDE model trained on relative abundances of genera selected by the skewness and kurtosis filtering outperforms all other models described above. The resultant model performs reasonably well on the test set (mean test set ROC-AUC = 0.82, accuracy = 74.5% precision = 0.65, recall = 0.71, F1-Score = 0.71). The results are presented in figure 6. Albeit not on the same validation dataset, our approach describes better results than the best submission in the DREAM challenge for term vs preterm prediction (ROC-AUC = 0.68, accuracy = 67%, sensitivity = 0.48, specificity = 0.79) [26], despite using microbial abundances only up to the end of the 2^nd^ trimester, i.e., the 24^th^ week of gestation, as opposed to the 32^nd^ week of gestation in the DREAM challenge.

**Figure 6:**
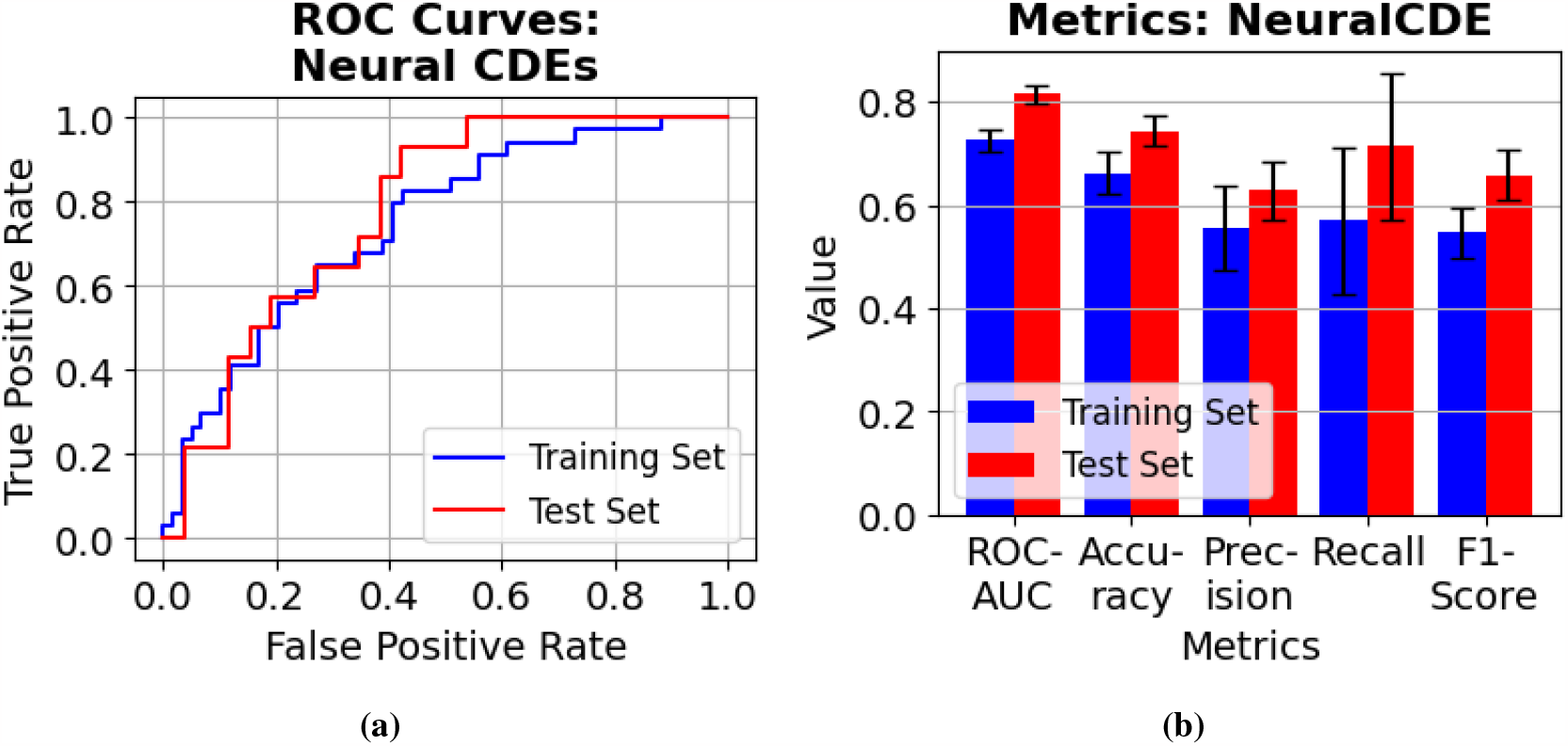
a ROC curves and b classification metrics on the training and test datasets for the Neural CDE model.

## 4 Discussion

Current *in vitro* or *in vivo* approaches lack the ability to detect PTB incidences at earlier stages reduces the effectiveness of prophylactic or therapeutic interventions that can be administered to mitigate neonatal health concerns associated with it. Current risk assessment approaches involve physical examinations, and factors such as cervical length may be used to estimate the risk. Additionally, abnormality in levels of biochemical markers such as Pregnancy-Associated Protein A (PAPP-A) [57, 29], Cervicovaginal Interleukins [42, 49], etc., may help detect PTB as well. However, there is a lack of definitive confidence intervals for these physical or biochemical tests. Recent studies have highlighted the utility of diversity of vaginal microbial communities towards PTB prediction, by establishing correlations between alpha-diversity indices associated with abundances of various microbial species (or genera) and incidences of PTB [18, 34, 31]. However, microbial communities are highly diverse across various individuals, and more so for individuals belonging to different ethnic populations [60, 30]. We have demonstrated above, that in a dataset with microbial profiles derived from ethnically and racially diverse subjects, diversity metrics could not accurately estimate PTB risk, and they perform worse than random predictors. This indicates that previously reported success of diversity indices in identifying subjects at high risk of PTB may be dataset dependent, either with respect to the subject cohort or to the pipeline used in computation of microbial abundance from 16S rRNA sequences, based on our observations on a mixed-race dataset with 16S rRNA sequences transformed using a standardized method.

Traditional ML methods, which have been explored previously in the context of predicting PTB using vaginal microbial species abundance, fail to learn an abundance signature associated with PTB in our dataset. As we have demonstrated above, vaginal microbial communities evolve throughout the duration of pregnancy, and abundance levels of certain species or genera may change significantly as the pregnancy progresses. Learning a PTB-associated signature in an evolving microbiome may be out of scope of such models as they are not designed to handle time series datasets. We explored the utility of LSTM, a type of RNN, which is able to work with sequential datasets. The architecture of RNN-based approaches requires input datasets to be regularly and continuously sampled. As far as human patient subject data is concerned, obtaining such a dataset is a challenge, as study subjects may not be regular or consistent in clinical visits. For this purpose, we filtered the dataset such that a single sample was present across each of the gestational weeks, which constituted the time intervals for LSTM. We suitably modified the LSTM workflow to overcome missing time intervals, however it proved to be incapable of learning any signature associated with PTB.

Neural differential equations have recently gained traction with regards to analyzing sequential data. Since it uses differential equations to model the temporal dynamics, it can handle irregularly and/or inconsistently sampled data. On top of that, neural CDEs are even capable of working with partially sampled datasets, and are more efficient than neural ODEs [36]. We found that neural CDEs were able to predict PTB incidences in our dataset with substantial accuracy, and outperformed all other approaches that we tested. To the best of our knowledge, this is the first effort towards modelling the temporal dynamics of vaginal microbial communities using deep learning, and the first instance of applying neural differential equations for a problem of this kind.

The DREAM challenge for PTB prediction[26], was issued in 2019 with the goal of driving efforts for PTB prediction using the vaginal microbiome. One of the sub-problems for the DREAM challenge consisted of predicting term births (>= 37 weeks of gestation) and preterm births (< 37 weeks of gestation) using vaginal microbiomes. The dataset for this challenge was derived from 9 different studies, and amounted to 3578 samples collected from 1268 individuals [26]. The dataset used in this study was also part of the DREAM challenge. We also considered using some of the other datasets in the DREAM challenge while outlining this study, but dropped either due to not being labelled week-wise or due to insufficient week-wise samples per patient, for modelling the temporal dynamics. Most of the top submissions in the challenge used tree-based classifiers. On our test dataset, neural CDEs show better predictivity (mean test set ROC-AUC = 0.82, accuracy = 75%, sensitivity = 0.71, specificity = 0.85) than the best submission in the DREAM challenge on their validation dataset (ROC-AUC = 0.69, accuracy = 67%, sensitivity = 0.48, specificity = 0.79) [26], despite using microbial abundances only up to the end of the 2^nd^ trimester, i.e., the 24^th^ week of gestation. The DREAM challenge for PTB prediction used taxonomic abundance data upto the 32^nd^ week of gestation. Our emphasis was on early-stage PTB prediction, and we were able to achieve better predictive performance in spite of restricting the input data till the 2^nd^ trimester.

Predictive approaches using data other than microbial communities also exist. For instance, Tarca et al. [61] report the results of the DREAM challenge for PTB prediction using the maternal blood transcriptome and the proteome. The top performing models report better results than what was reported on microbial communities, with a ROC-AUC of 0.76 when proteomics data from weeks 27-33 was used. However, in another sub-challenge where early-stage data (weeks 17-22) was used, the top performing model had a ROC-AUC of 0.62. Obtaining blood transcriptomic or proteomic data may pose difficulties due to the involvement of invasive procedures requiring clinical expertise to perform. On the other hand, microbial abundance data is sourced from vaginal swabs, which can be obtained without invasive procedures, by patient subjects themselves. Several attempts have been made at predicting PTB using biochemical marker [2, 38], however, obtaining such data may require regular clinical visits, and their viability in racially-diverse populations is unknown.

Poorer and remote parts of the world may even lack the medical infrastructure or presence of adequate facilities that are required for PTB assessment and prevention. For example, in remote areas in India, there are clinics called “Anganwadis”, which roughly translates to “courtyard shelter”. As of 2018, the Ministry of Women and Child Development reports the existence of 1.4 million Anganwadi centres spread out across the country. Anganwadis provide limited healthcare facilities for maternal and infant health and lack the funding and facilities, or even trained medical personnel required to mitigate PTB and its ill-effects. Given the simplicity of obtaining samples from which microbial abundance is derived, reliable approaches for PTB risk assessments developed on microbiota will greatly help such remote clinics.

## 5 Limitations and Future Work

While we have demonstrated the capability of Neural CDEs towards PTB prediction using the vaginal microbiome, further effort can be made for increasing its clinical viability. Firstly, our dataset is limited in size (133 patient subjects), and we believe that larger datasets with better racial and ethnic representation may help learn signatures which take into account the diversity of vaginal microbiomes across individuals/races. Secondly, predicting the extent of preterm birth (extremely preterm, very preterm, moderate to late preterm) is also important as far as administering interventions is concerned, as they may have varying impact on maternal and infant health and may require different strategies. This may be achieved by predicting the gestational week of delivery, or by treating PTB as a multi-class problem with different extents of PTB as the classes, on more high-quality datasets. We strongly believe that vaginal microbial communities may be the key to achieving early-stage PTB prediction, and our findings strongly encourage future efforts for pregnancy microbiome data generation and further refinements in modelling procedures, which may take us closer to achieving full clinical viability.

## 6 Acknowledgement

The authors of this study declare that they are employees of Tata Consultancy Services Limited - a commercial company. The company has provided support for this study in the form of salaries to authors, but did not have any additional role in the study design, data collection and analysis, decision to publish, or preparation of the manuscript. The authors would like to thank Dr. Mohammed Haque, Dr. Anirban Dutta, and Mr. Nishal Kumar Pinna for their help in processing the 16S rRNA sequences from the SRA cloud, and Mr. Sunil Nagpal and Dr. Anirban Dutta for outlining the state of the art for PTB prediction using diversity metrics. We would also like to thank Dr. Rajgopal Srinivasan for proofreading and helping improve the manuscript.

## 7 Author Contribution

M.B. and K.K. designed the study. K.K. performed the computational analyses along with assistance from M.B., and M.B. and K.K drafted the final manuscript.

