## Supplementary Material for "Neural differential equations enable early-stage prediction of preterm birth using vaginal microbiota"

### Deep Learning Methods for Early-Stage Prediction of Preterm Birth Using Vaginal Microbiota

Supplementary Material

#### 1 Machine Learning Classifiers: Hyperparameters

##### 1.1 Hyperparameter search spaces

Hyperparameters for both, the Random Forest (RF) and Decision Tree (DT) models were tuned by carrying out 3-fold cross-validation on the training set. For each set of hyperparameters, the training set was divided into 3 folds, and classification performance was evaluated on each fold after training on the 2 remaining folds. Accuracy, ROC-AUC, precision, recall and F1-score was computed for each fold. Finally, the optimal hyperparameters were selected based on mean F1-score (i.e., the hyperparameter set with the highest mean F1-score was selected as the optimal set). The search spaces of the hyperparameters are presented in supplementary table 1. The explanation and significance of parameters can be found in the scikit-learn documentation.

| Parameter | Search Spaces |
|---|---|
| n_estimators (RF only) | 50, 100, 150 |
| max_depth | None, 2, 3, 4 |
| min_samples_split | 0.25, 0.33, 0.5, 2, 4 |
| max_features | sqrt, log2, 0.25, 0.33, 0.5, 2, 4 |
| max_leaf_nodes | None, 2, 4 |
| min_samples_leaf | 0.25, 0.33, 0.5, 2, 4 |
| criterion | entropy, gini, log_loss |
| splitter (DT only) | best, random |
| class_weight | balanced, balanced_subsample |

Supplementary Table 1: Hyperparameter search spaces for the Random Forest and Decision Tree classifiers. The search spaces for both models are common unless specified otherwise.

##### 1.2 Optimal Hyperparameter Values

The optimal values of the hyperparameters, as identified by grid search, are listed in supplementary table 2.

| Parameter | Optimal Value | |
|---|---|---|
| | Random Forest | Decision Tree |
| n_estimators | 50 | - |
| max_depth | 4 | 3 |
| min_samples_split | 0.33 | 0.5 |
| max_features | 0.25 | 0.25 |
| max_leaf_nodes | 2 | 4 |
| min_samples_leaf | 0.33 | 0.2 |
| criterion | entropy | gini |
| splitter | - | best |
| class_weight | balanced_subsample | balanced |

Supplementary Table 2: Optimal hyperparameter values for the Random Forest and Decision Tree classifiers.

**2. Long Short-Term Memory (LSTM):**

**2.1 Implementation details**

To facilitate the LSTM model to make predictions on a non-continuous and irregularly sampled dataset, we made two modifications to it. Let $w_0$ be the first time step for which data is available. Firstly, the hidden state is initialized by an embedding layer, which takes $w_0$ as input, and the cell state is initialized as a zero vector. Since week 9 is the earliest possible time point for which data is used, we denote it as t = 0. The LSTM model is run from weeks 9 to 24 (t = 0 to t = 15). From t = 0 to t = $w_0$ - 1, the LSTM cells at each time step do not update the hidden state, and simply act as dummy cells. At t = $w_0$, the hidden state is still the same as what was initialized with the embedding, and the LSTM cell at t = $w_0$ updates the hidden state based on the input data.

The LSTM cell at each time step is followed by a linear layer which generates the "forecast", or the predicted genera abundance at that time step. As part of our second modification, for the time steps beyond t = $w_0$ for which input data is missing, we use this forecast to update the hidden state. Since the missing sampling intervals are different for each subject, we have to process each subject with its own set of rules to update the model weights. Thus, the batch size is restricted to 1.

**2.2 Model Parameters**

We used the Adam optimizer implemented within pytorch for training the model. The model parameters for LSTM are listed in supplementary table 3.

| Parameter | Value | Explanation |
|---|---|---|
| Input channels | 30 | Dimension of the input (number of genera in the input dataset) |
| Hidden channels | 128 | Dimension of the hidden state |
| Embedding input dimensions | 14 | Number of possible inputs to embedding layer (i.e., number of unique weeks between 9 and 24 which serve as first available data sample) |
| Learning rate | 1e-4 | Learning rate for updating weights |
| Learning rate decay | 1 (No decay) | Reduction in learning rate across epochs |
| Loss function | Binary cross-entropy | The function to minimize while optimizing the LSTM model |
| Number of epochs | 35 | Number of epochs (iterations) for which the model is trained |
| Batch size | 1 (Fixed) | Number of samples per training batch. Batch size is restricted to 1 due to different time steps for which data is missing, which forces handling each sample individually |

Supplementary Table 3: Parameters for the long short-term memory model

**3. Neural Controlled Differential Equations (CDEs)**

**3.1 Training using adjoint backpropagation**

Neural differential equation models are solutions to a time-dependent system whose function is a neural network that can be learned. This neural network is typically a linear fully connected network, however, a non-linearity can be introduced by using a tanh or a relu activation. In case of CDEs, there exists a "hidden state" (h), essentially a parametrized version of the input data, and the ODE can be represented as:

$$\frac{dh}{dt} = f_\theta(h)\,\frac{dX}{dt}$$

Where $f_\theta$ is the neural network with parameters $\theta$. Training the model essentially refers to update these parameters based on the gradient of the cost function (J) with respect to the parameter vector, scaled by a learning rate ($\alpha$).

$$\theta_{i+1} = \theta_i + \alpha \frac{dJ}{d\theta}$$

And the gradient of the cost function can be represented as:

$$\frac{dJ}{d\theta} = \int_0^T g(h, \theta)\ dt$$

Where g is the loss function (binary cross-entropy in our case). The above expression essentially maps this gradient to a scalar, using which the parameters are updated. This can be further represented as:

$$\frac{dJ}{d\theta} = \int_0^T \frac{\partial g}{\partial \theta} + \frac{\partial g}{\partial h}\frac{dh}{d\theta} dt$$

Adjoint sensitivity analysis or the adjoint state method essentially refers to computing the cost function gradient using the above expression in an efficient manner. The exact mathematical derivation for this is comprehensively described in Chen et al. [1].

**3.2 Model parameters**

| Parameter | Value | Explanation |
|---|---|---|
| Input channels | 6 | Dimension of the input (number of genera in the input dataset) |
| Hidden channels | 64 | Dimension of the hidden state |
| Output channels | 1 (Fixed) | Dimension of the output (fixed to 1, since we are only outputting a single probability) |
| Interpolation type | Cubic | Method of interpolation of missing data from nearest observations |
| Learning rate | 1e-4 | Learning rate for updating weights |
| Learning rate decay | 0.5 | Reduction in learning rate across epochs |
| Learning rate decay step size | 15 | Number of epochs after which learning rate decay is applied |
| Loss function | Binary cross-entropy | The function to minimize while optimizing the LSTM model |
| Positive class weight | 1.5 | Weighted penalty to apply to negative class predictions (i.e., encourage predictions of positive class) |
| Batch size | 10 | Number of samples per training batch |

Supplementary Table 3: Parameters for the Neural CDE model.
